# Cross-fostering reveals that acoustic communication during parental care alleviates prenatal disadvantages in beetle offspring

**DOI:** 10.1101/2025.09.24.678224

**Authors:** Stella Mangold, Sandra Steiger, Taina Conrad

**Author notes:** **Corresponding author.** E-mail address (T. Conrad).

## Abstract

Acoustic communication plays an important role in social interactions for many animals. It is used in a broad range of behaviours, such as parental care. Acoustic communication is also involved in the brood care of *Nicrophorus vespilloides* (HERBST 1783), which, together with the other burying beetles, are one of the rare exceptions among insects in that they practice biparental brood care. Previous studies showed that a lack of acoustic communication in burying beetles negatively affects the weight and survival rate of their offspring (Conrad et al. 2024). However, it remained unclear whether these results were mainly based on prenatal or postnatal effects of silencing. Therefore, in this study we wanted to find out whether negative effects on *N. vespilloides* larvae, which are due to a lack of prenatal acoustic communication between the parents, can be compensated by the postnatal brood care of stridulating foster parents.

For this purpose, we applied parafilm on the pars stridens of some beetle pairs so that they could no longer stridulate. In a 2×2 cross-fostering experiment, larvae originating from silenced or stridulating parents were reared by either silenced or stridulating foster parents. The larvae of silenced parents showed a lower hatching weight than the larvae of control parents. At dispersal, larvae with stridulating original and foster parents were significantly heavier than those reared by silenced foster parents, from either origin. At dispersal, larvae with stridulating original and foster parents were significantly heavier than those reared by silenced foster parents, from either origin. Thus, the stridulating foster parents were apparently able to partially compensate the initial weight deficits of the larvae from silenced original parents. There were no effects of silencing on offspring survival.

Our experiment demonstrates that post-hatching acoustic communication during brood care of *N. vespilloides* is of greater importance for larval growth than pre-hatching acoustic communication. Still, the inability to stridulate compromises the quality of both prenatal and postnatal care. It is conceivable that the acoustic communication of the parents provides orientation for the larvae when feeding, or that the adults coordinate their tasks and negotiate their parental investment. Our findings contribute further to our understanding of the function of parental communication during brood care.

## Introduction

Family life is a very fundamental form of sociality and thought to be the first step in the evolution to a more complex social life (Szathmáry and Maynard Smith 1995; Kramer and Meunier 2018). Parental care in turn encompasses all activities and characteristics of parents that contribute to the increase of their offspring’s fitness and are likely to have evolved for this purpose or are currently maintained for this reason (Royle et al. 2012a). Brood care can be divided into prenatal and postnatal stages. For example, the construction of nests is usually a form of prenatal brood care, as are the nutrients and resources parents supply within eggs, while the feeding of hatched or born offspring is a form of postnatal brood care. Activities such as temperature regulation and guarding can fall into both categories. Every parental investment that increases the fitness of an offspring inevitably leads to a decrease in investment in other offspring (Trivers 2017). This trade-off contributes to different strategies regarding the intensity and duration of parental care between species (Royle et al. 2012a). While biparental care is the most common form of parental care in birds (Cockburn 2006), it seldomly occurs in other taxa. This is also the case in insects, most of which do not engage in brood care (Alcock and Rubenstein 2019). Termites (Isoptera), wasps and ants (Hymenoptera) are generally known for their alloparental care, while biparental care is extremely rare (Hölldobler and Wilson 1990; Eggert and Müller 1997; Shreeves and Field 2007; Korb et al. 2012; Royle et al. 2012a; Field et al. 2020).

In families, communication with each other is needed to keep this social construct functional, as the interests of the individuals involved do not necessarily align (Royle et al. 2012b; Trivers 2017; Kramer and Meunier 2018; Prang et al. 2022). Although communication is a key component in family life and parental care, it can be difficult to understand the specific role certain signals play. One form of communication is acoustic communication, which has been shown to play an important role in the context of parental care and brood care in many species especially among vertebrates (Royle et al. 2012a; Boucaud et al. 2016; Moss et al. 2023). For example, a recent study in poison frogs was able to show that they have specific calls associated with trophic egg feeding, a cooperative parental behavior. These frogs therefore seem to use acoustic signals in the coordination of their parental tasks (Moss et al. 2023). Additionally, studies have looked at the effects of prenatal and postnatal acoustic communication, especially in birds (Rumpf and Tzschentke 2010; Mariette and Buchanan 2016; Mariette 2019). However, the function of acoustic communication in the context of brood care in insects remains largely unexplored and almost nothing is known about prenatal vs postnatal effects.

*Nicrophorus* beetles with their biparental care behavior are ideal model organisms to investigate acoustic communication during brood care further. These beetles are known for their family life, which has been extensively studied and observed (Pukowski 1933; Eggert and Müller 1997; Scott 1998; Capodeanu-Nägler 2018; Potticary et al. 2024). The reproduction of *Nicrophorus* beetles occurs on a small vertebrate carcass. The parents bury the cadaver in the soil and shape it into a ball (Pukowski 1933). As the offspring grow, the parents defend the carcass and maintain its integrity. A unique aspect of postnatal brood care in *Nicrophorus* beetles is the direct feeding of begging offspring with pre-digested, regurgitated food by the parents (Eggert and Müller 1997; Scott 1998). After an initial time period when larvae rely heavily on their parents for feeding, the larvae are fully capable of feeding themselves and feeding by parents becomes less important over time (Smiseth et al. 2003; Capodeanu-Nägler et al. 2016). Meanwhile both parents produce stridulations throughout the entire brood care. (Pukowski 1933; Niemitz and Krampe 1971; Niemitz 1972; Niemitz and Krampe 1972; Scott 1998; Hall et al. 2013). Like many other insect species, the stridulation apparatus of *Nicrophorus* beetles consists of two pars stridens (scraping ridges), and two plectra (scraping edges) located on the fourth (female) or fifth (male) abdominal segment and on the underside of the elytra. Sounds are produced by rubbing a plectrum over a pars stridens (Pukowski 1933; Schumacher 1973; Hall et al. 2013; Schrader and Galanek 2022). There are fine differences in the morphology of the stridulation apparatus and the sound spectrum between individual *Nicrophorus* species (Hall et al. 2013). It has already been shown that *Nicrophorus* beetles stridulate during reproduction, such as during mating or brood care, and that this serves communication (Phillips et al; Niemitz and Krampe 1972; Huerta et al. 1992; Hall et al. 2013; Hall et al. 2015; Conrad et al. 2024).

In a previous study we used burying beetles to show the impact acoustic communication has on parental care. We were able to show that offspring of silenced parents fare worse than their control counterparts (Conrad et al. 2024). In *Nicrophorus vespilloides*, this disadvantage already begins pre-hatching with the size of the eggs, as silenced parents produce smaller eggs and consequently smaller larvae. Although the larvae are capable of increasing their larval weight to catch up to the weight of control larvae, their survival rate is significantly lower (Conrad et al. 2024). However, it remains unclear whether this is due to the initial disadvantage at hatching or due to impaired post-hatching care. So, does the silencing of the parents have only prenatal effects in *Nicrophorus vespilloides* or are there also postnatal effects due to lesser quality care of the parents? In order to better understand this intricate relationship between parents and offspring, particularly the importance of their communication, we used a full factorial cross-fostering experiment design. Newly hatched larvae from silenced or stridulating parents were reared either by silenced or by stridulating foster parents, allowing us to separate the effects of pre-hatching from post-hatching care.

## Materials and Methods

### Rearing and maintenance of beetles

The beetles that were used for the experiment were descendants of beetles collected from carrion-baited pitfall traps. *N. vespilloides* were caught in a forest near Bayreuth, Germany (49°55’18.192’’N, 11°34’19.9488’’E). All beetles were maintained in temperature-controlled chambers at 20 °C on a 16:8 h light:dark cycle. Before the experiments, groups of up to 5 adults of the same sex and family were kept in small plastic containers (10 × 10 cm and 6 cm high) filled with moist coconut coir. Beetles were fed whole fly larvae (*Lucilia sericata*) ad libitum twice a week. At the time of our experiments, beetles were virgin and between 20 and 30 days of age.

### Mating pairs and silencing of beetles

Mating pairs were chosen randomly from first generation lab populations, their pronotum width documented with a stereo microscope equipped with a camera (Stemi 305, Zeiss, Berlin, Germany) and then assigned randomly to the silenced or control group similar to Conrad et al (2024). Beetles were then anesthetized using CO_2_ and consequently silenced by gluing a small (approx. 4mm”) piece of parafilm (Bernis Inc., Neenah, Wisconsin, USA) onto the stridulatory organ using super glue (Super Glue Ultra Gel, Pattex ©, Henkel AG &Co KGaA, Düsseldorf, Germany). The control beetles were treated the same way but the parafilm was placed onto the lower part of the abdomen where it would not interfere with the stridulatory organ. After the attachment of the parafilm, beetles were kept anesthetized for approximately 10 more minutes to allow the glue to fully dry. Successful silencing was checked throughout the experiment visually as well as audibly while handling.

### Experimental design

To disentangle the effects of parental acoustic communication during pre-hatching and post-hatching brood care on offspring fitness, we used a full factorial design with four groups: silenced parents rearing newly hatched larvae from silenced parents, silenced parents rearing newly hatched larvae from control parents, control parents rearing newly hatched larvae from silenced parents and control parents rearing newly hatched larvae from control parents.

Reproduction was induced by providing each mating pair with a 20 g (± 2.5 g) thawed mouse carcass (Frostfutter.de—B.A.F Group GmbH, Germany). Since *N. vespilloides* is active during daylight, mice were provided in light, and beetles were moved to the dark after 5 h when the carcass was buried. After the egg-laying period but before larvae hatched (Capodeanu-Nägler et al. 2016; Conrad et al. 2024), parents and the carcass were transferred to new plastic containers filled with coconut coir. The eggs were left to hatch in the old container, which we checked every 4 h for the presence of newly hatched larvae. We weighed the larvae when they hatched (0 h), before providing each couple of beetles with a brood of 10 newly hatched larvae of mixed parentage (within either group of silenced or control beetles) to control for variation between families and individual differences in behavior (Rauter and Moore 1999).

Females exhibit temporally-based kin discrimination, during which they kill any larvae arriving on the carcass before their own eggs would have hatched but accept larvae that arrive after their own eggs have begun to hatch (Müller and Eggert 1990). Therefore, we provided couples with larvae only after their own larvae had begun hatching. We established broods to attain a minimum sample size of 18 for each group (n = 18 silenced parents + larvae from silenced parents, n = 26 silenced parents + larvae from control parents, n = 28 control parents + larvae from silenced parents and n = 24 control parents + larvae from control parents).

As larval begging and parental feeding is most pronounced in the first 48h (Smiseth et al. 2003; Capodeanu-Nägler et al. 2016), larvae were weighed again after 48h and at dispersal, at which points survival was noted each time. After dispersal larvae were allowed to pupate and after approximately 30 days newly emerged adults were counted and sexed.

### Statistics

For all analysed response variables, we fitted fixed-effects models with an interaction term for treatment parents (either silenced or control) * origin of larvae (either from silenced or control parents) and size of male parent * size of female parent. We also added carcass weight as fixed effects:

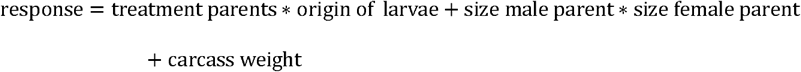

Generalized linear models (GLMs) with Gaussian distributed error structure and log link were fitted to continuous response variables (average larval weight 48 hours after hatching, average larval weight at dispersal, growth rate till 48 hours ((avg. weight at 48 h – avg. weight at 0 h) /avg. weight at 0 h, growth rate 48 hours till dispersal ((avg. weight at dispersal – avg. weight at 48 h) /avg. weight at 48 h). Generalized linear models (GLMs) with Poisson distributed error structure and log link were fitted to survival counts and number of new adults.

Residuals of linear models were checked visually based on standard residual plots and by plotting residuals against predictors. Residuals of GLMs were checked using *DHARMa*, version 0.4.6 (Hartig, 2017). The contributions of different predictors to the variance in the data were tested via type II ANOVAs and likelihood ratio tests (GLMs) using the Anova() function, *car* package (Fox & Weisberg, 2019). All analyses were done in R version 4.3.1. All graphs were produced using Sigma Plot 14.0 (Systat Software, Chicago, IL, U.S.A.).

## Results

At hatching, just as in our previous study, the origin of the larvae had a significant effect on the weight of the larvae - with those of control parents being heavier (GLM, df = 1, F_larvae_ = 18.79, P < 0.001; Fig. 1). None of the other factors had an effect (GLM, df = 1, P > 0.05).

**Figure 1:**
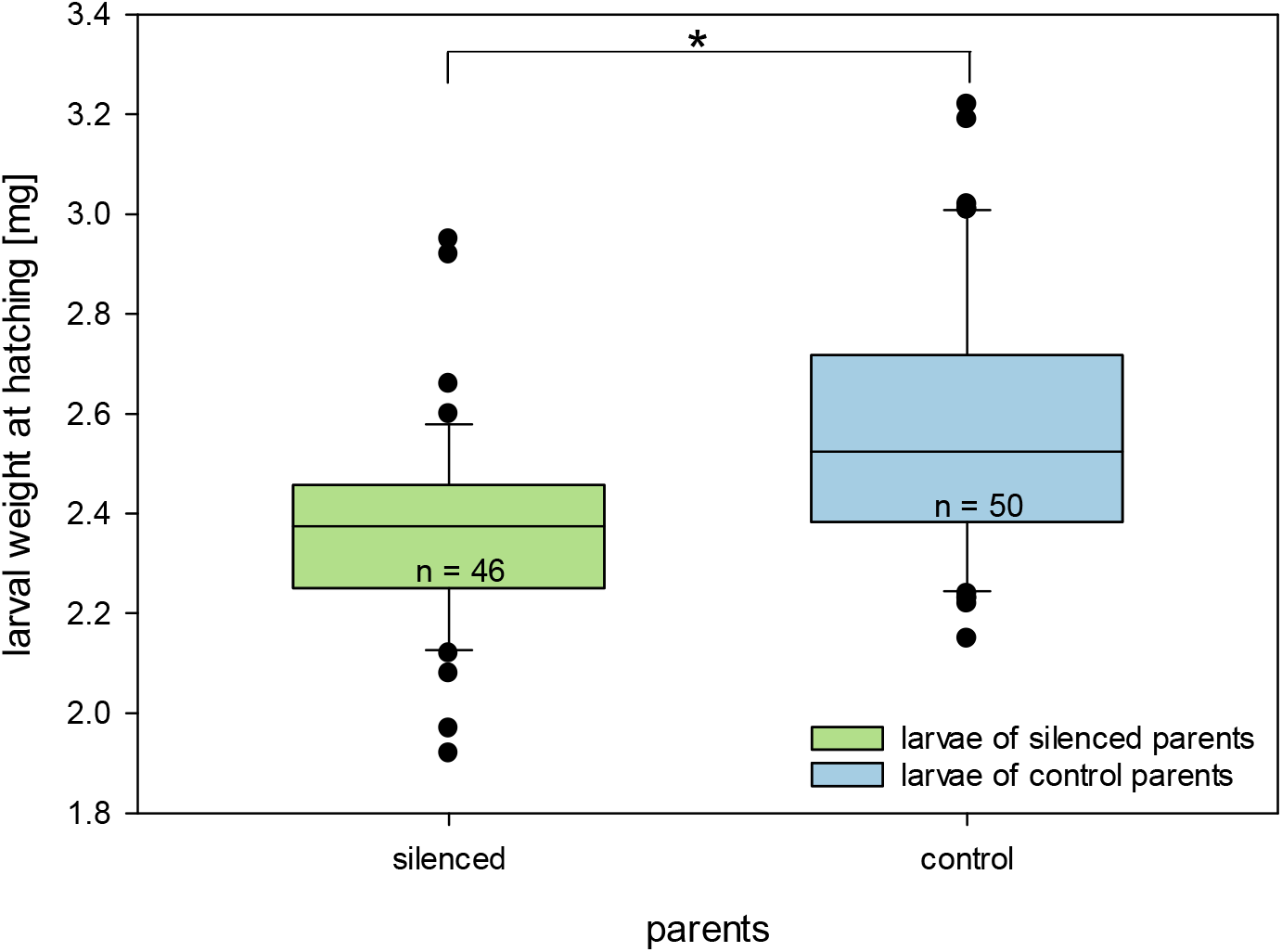
Larval weight at hatching of silenced and control parents. The numbers within the box represent the sample size (n). The medians, quartiles and outliers (data points beyond 1.5×IQR from the quartiles) are shown. Whiskers represent the data range without outliers. Significant differences are marked by stars (GLM with subsequent Anova and pairwise Tukey test, *P < 0.05).

After 48 hours, we found no difference in larval weight between the groups (GLM, F_parents_ = 0.05, p = 0.83; F_larvae_ = 0.46, p = 0.50; Fig. 2). However, growth rate until 48 hours was significantly affected by the origin of the larvae (GLM, df = 1, F_parents_ = 0.09, p >0.05, F_larvae_ = 4.90, p = 0.03) with the growth rate being higher in larvae from silenced parents. In contrast, growth rate between 48 hours and dispersal was significantly affected by the treatment of the foster parents (GLM, F_parents_ = 4.24, p = 0.04, F_larvae_ = 0.11, p > 0.05) with the growth rate being lower in silenced groups (Fig.3).

**Figure 2:**
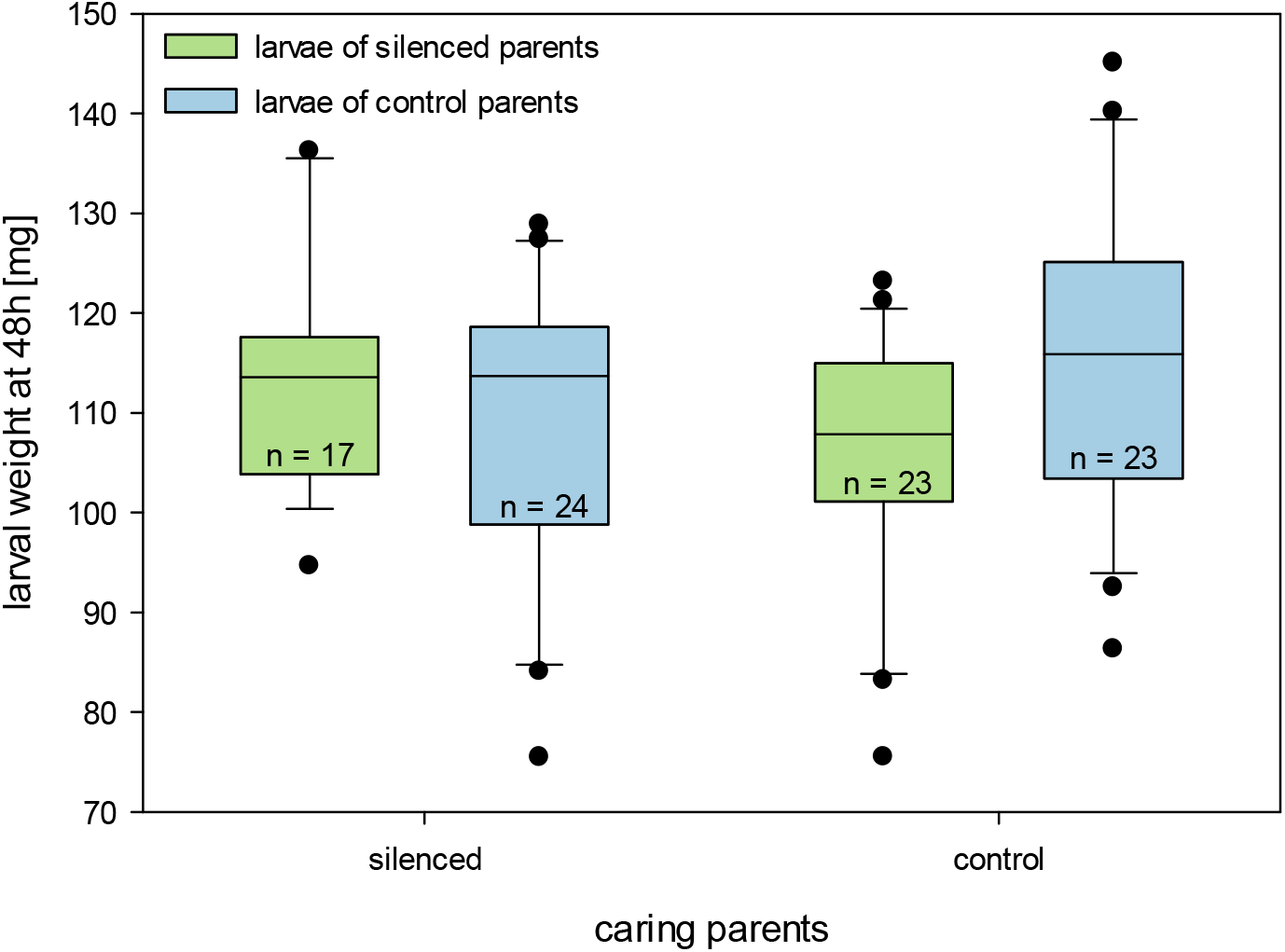
Larval weight at 48h of silenced and control parents. The numbers within the box represent the sample size (n). The medians, quartiles and outliers (circles) are shown. There were no significant differences (GLM with subsequent Anova and pairwise Tukey test, P > 0.05).

**Figure 3:**
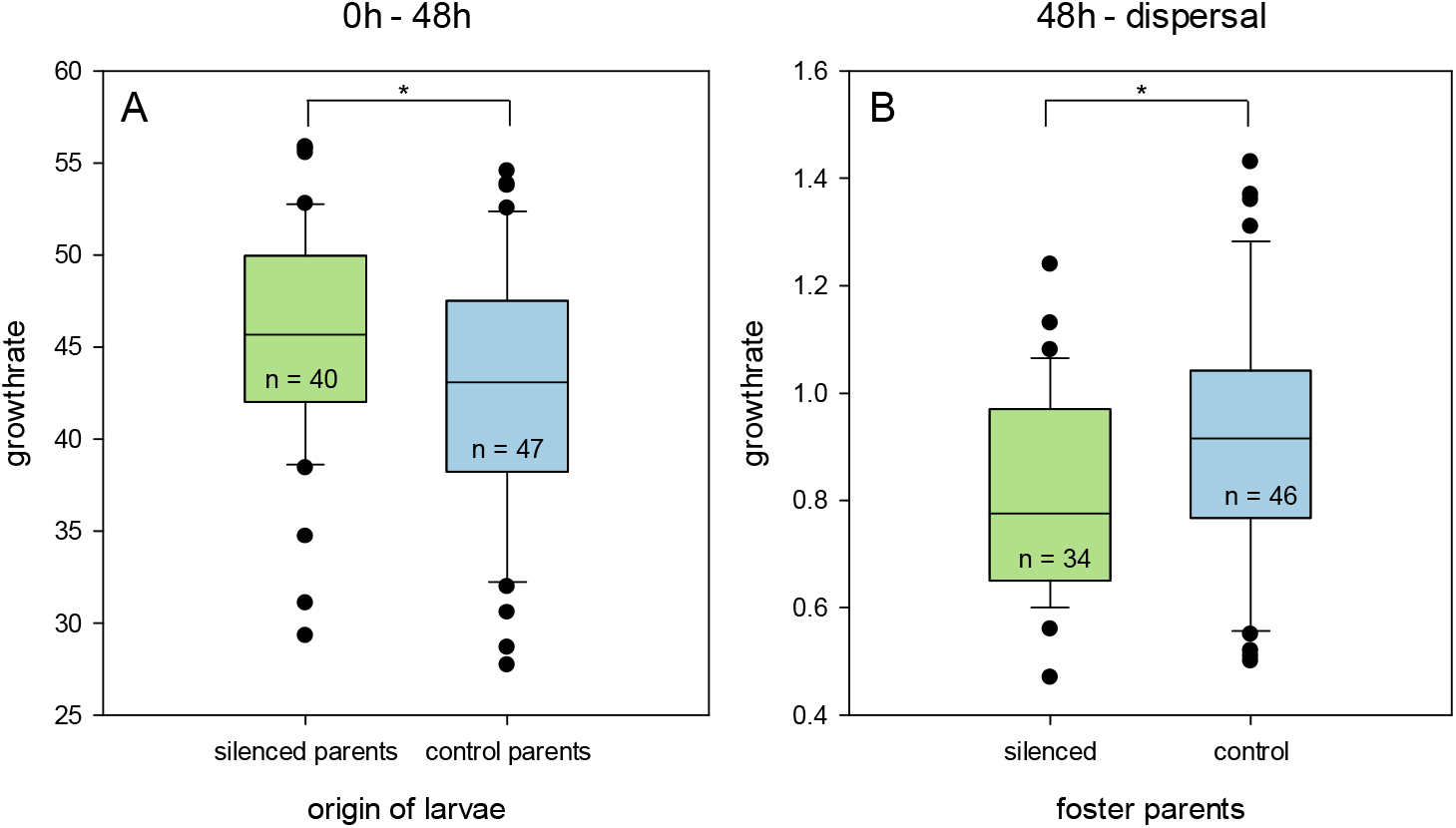
Growth rate of larvae from silenced or control parents 0h till 48h (A) and 48h till Dispersal (B) based on either the “origin of the larvae” (A) or the “treatment of the foster parents” (B) The numbers within the box represent the sample size (n). The medians, quartiles and outliers (data points beyond 1.5×IQR from the quartiles) are shown. Whiskers represent the data range without outliers. (GLM with subsequent Anova and pairwise Tukey test, Tukey,*P < 0.05)

At dispersal we found that treatment of the parents had a significant effect on larval weight while the origin of the larvae had no effect on the weight (GLM, df = 1, F_parents_ = 15.64, p < 0.001; F_larvae_ = 1.73, p = 0.19; Fig. 4). The larvae of silenced parents cared for by silenced parents had the lowest average weight and differed significantly from larvae of control parents cared for by control parents (GLM, Tukey, p < 0.01). Additionally, the larvae of control parents cared for by control parents weighed significantly more than larvae of control parents cared for by silenced parents (GLM, Tukey, p < 0.01). However, the larvae from silenced parents that were cared for by control parents did not differ significantly from either group (GLM, Tukey, p > 0.05)

**Figure 4:**
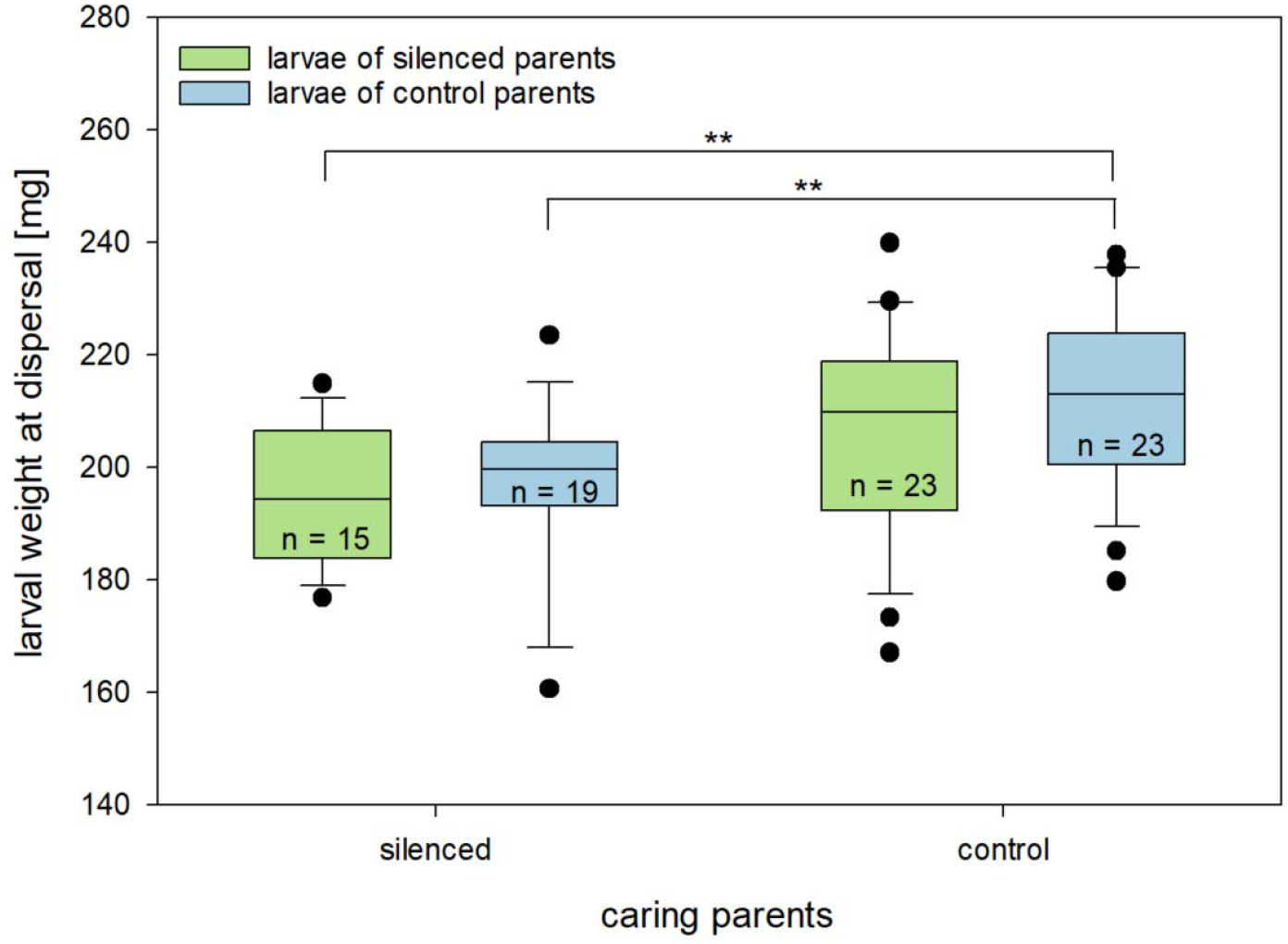
Larval weight at dispersal of larvae cared for by silenced as well as control parents. The numbers within the box represent the sample size (n). The medians, quartiles and outliers (circles) are shown. Significant differences are marked by stars (GLM with subsequent Anova and pairwise Tukey test, Tukey, **P < 0.01).

Additionally, the size of the female influenced larval weight with larger females having slightly smaller larvae (GLM, df = 1, F_SizeFemale_ = 4.6679, P < 0.05) – none of the other factors had any effect (GLM, df = 1, P > 0.05).

There was no significant difference in larval survival between any of the four groups at any time point (GLM, df = 1; Χ^2^_48hparents_ = 2.12; Χ^2^_48hlarvae_ = 0.29; Χ^2^ _dispersalparents_ = 3.02; Χ^2^ _dispersallarvae_ = 0.43, p > 0.05). On average, 2-3 larvae died during brood care. There was also no difference between the groups regarding the number of newly hatched adults - neither for females (GLM, df = 1; Χ^2^_parents_ = 1.86; Χ^2^_larvae_ = 0.19, p > 0.05) nor for males (GLM, df = 1; Χ^2^_parents_ = 1.68; Χ^2^_larvae_ = 0.18, p > 0.05).

## Discussion

Extending on our previous study we were able to show that the initial ability of larvae of silenced parents to “catch up” to their control counterpart seems to be driven by the status of the larvae at birth. Growth rate in the first 48 h is not affected by the status of foster parents and is higher in larvae from silenced parents, which were originally smaller. However, after 48 h, when larval weight has equalized over all groups, growth rate is affected by the status of the foster parents: larvae which were cared for by silenced parents have lower growth rates, leading to the differences in larval weight we find at dispersal. This means that silencing reduces the effectiveness of post-hatching care. Furthermore, the fact that the larvae from silenced parents, which were raised by control parents, did not differ significantly in weight from control larvae raised by control parents puts further emphasis on the idea that control parents can care better than silenced parents can.

Our results are quite surprising because we would have expected the effect of parental treatment to be strongest in the first 48 h after hatching when, according to literature, the dependence on brood care is the strongest and parental feeding is highest (Smiseth et al. 2003; Capodeanu-Nägler et al. 2016). It is possible that during the first 48 hours, when feeding is happening more often, communication is not as important, since tasks do not need to be allocated as much if one parent is already feeding most of the time. It is also possible that the lower weight at hatching leads to a significant increase in begging behavior in those larvae and consequently more feeding from the parents resulting in a higher initial growth rate. Other studies have shown that begging behavior can be a strong driver of increased feeding behavior and that begging behavior in *Nicrophorus* is both tactile but also chemical (Rauter and Moore 1999; Smiseth et al. 2003; Sahm et al. 2024).

There have been many studies in birds showing that acoustic communication of parents has a positive effect on the offspring (Magrath et al. 2010; Boucaud et al. 2017; Mariette 2019). For example, in black-backed gulls, *Larus fuscus*, pairs contributing more equally to incubation vocalized more and had a higher reproductive output (Kavelaars et al. 2019). In turn, the disruption of vocalizations can have detrimental effects. This is the case of blue tits, where playbacks of road noise led to a disrupted vocal communication and subsequent reduced parental provisioning (Lucass et al. 2016).

We believe that there are two possible reasons for the reduction in quality of care from silenced parents. First, acoustic communication between parents facilitates the coordination of their tasks. Such coordination can increase the efficiency of parental care and ultimately lead to increased reproductive success, as has been shown in zebra finches, *Taeniopygia guttata*, which also exhibit biparental care (Mariette and Griffith 2012, 2015). Burying beetles show a very high ability for task allocation, such as offspring feeding, carcass maintenance and carcass protection. Previous work has shown a negative relationship between female and male feeding, with males increasing their feeding when females reduce theirs. (Fetherston et al. 1990; Smiseth and Moore 2004; Lambert and Smiseth 2024). This pattern implies coordination between partners, which could be mediated through acoustic signaling. Without the ability to coordinate their tasks through acoustic communication, parental care may be less efficient, and the larvae may therefore grow more slowly and reach a smaller size.

The second possibility is that without the ability to stridulate, communication between parents and their larvae may be disrupted. Although larvae can find their way to the food source through olfactory cues (Pukowski 1933), it is possible that parental calls provide additional orientation (Niemitz and Krampe 1972). It is also possible that the offspring orient themselves to the parents’ calls during feeding, as is the case in many birds (Clay et al. 2012), and therefore silencing the parents may have a negative impact on the growth of the larvae. This is supported by observations from other studies, during which the parent was seen stridulating atop the carcass with the larvae responding by coming up for feeding (Prang, Schückel, Müller pers. comm.; Conrad, pers. obs.). This behavior seems to be disrupted significantly by silencing the parents (Conrad, per. obs.).

In conclusion, our study demonstrates that both prenatal and postnatal acoustic communication contribute to offspring fitness in burying beetles, paralleling findings in many bird species. This emphasizes the important role of acoustic communication during family life. Future studies should look further into the importance of maternal versus paternal communication. By only silencing one partner it would be possible to show if female or male communication drives the prenatal or postnatal disadvantages.

## Acknowledgements

We would like to thank Magdalena Mair for valuable statistical advice and Markus Conrad for advice on the manuscript. The research was supported by a grant of the German Research Foundation (DFG) to T.C (CO 2181/1-1 Project number: 424437154).

## CRediT

Stella Mangold (formal analysis, investigation, visualization, writing – original draft); Sandra Steiger (writing – review &editing, resources); Taina Conrad (conceptualization, formal analysis, funding acquisition, methodology, resources, supervision, visualization, writing – review & editing)

## References

Alcock J, Rubenstein DR. 2019. Animal behavior. Eleventh edition. Sunderland Massachusetts: Oxford University Press.

Boucaud IC, Mariette MM, Villain AS, Vignal C. 2016. Vocal negotiation over parental care? Acoustic communication at the nest predicts partners’ incubation share. Biological Journal of the Linnean Society:322–336.

Boucaud IC, Perez EC, Ramos LS, Griffith SC, Vignal C. 2017. Acoustic communication in zebra finches signals when mates will take turns with parental duties. Behavioral Ecology. 28:645–656.

Capodeanu-Nägler A. 2018. Parental care and variation in offspring dependency in burying beetles of the genus Nicrophorus. Dissertation. Ulm: University of Ulm, Institute of Evolutionary Ecology and Conservation Genomics. eng.

Capodeanu-Nägler A, Keppner EM, Vogel H, Ayasse M, Eggert A-K, Sakaluk SK, Steiger S. 2016. From facultative to obligatory parental care: Interspecific variation in offspring dependency on post-hatching care in burying beetles. Sci Rep. 6:369.

Clay Z, Smith CL, Blumstein DT. 2012. Food-associated vocalizations in mammals and birds: what do these calls really mean? Animal Behaviour. 83:323–330.

Cockburn A. 2006. Prevalence of different modes of parental care in birds.

Conrad T, Mair MM, Müller J, Richter P, Schödel S, Wezstein A-K, Steiger S. 2024. The impact of acoustic signalling on offspring performance varies among three biparentally caring species. Animal Behaviour. 217:13–20.

Eggert A-K, Müller JK. 1997. Biparental care and social evoliution in burying beetles: lessons from the larder. In: Crespi BJ, Choe JC, editors. The evolution of social behavior in insects and arachnids. Cambridge: Cambridge University Press.

Fetherston IA, Scott MP, Traniello JFA. 1990. Parental Care in Burying Beetles: The Organization of Male and Female Brood-care Behavior. Ethology. 85:177–190.

Field J, Gonzalez-Voyer A, Boulton RA. 2020. The evolution of parental care strategies in subsocial wasps. Behavioral Ecology and Sociobiology. 74:78.

Hall CL, Howard DR, Smith RJ, Mason AC. 2015. Marking by elytral clip changes stridulatory characteristics and reduces reproduction in the American burying beetle, Nicrophorus americanus. J Insect Conserv. 19:155–162.

Hall CL, Mason AC, Howard DR, Padhi A, Smith RJ. 2013. Description of acoustic characters and stridulatory pars stridens of Nicrophorus (Coleoptera Silphidae): A comparison of eight North American species. Ann. Entom. Soc. Amer. 106:661–669.

Hölldobler B, Wilson EO. 1990. The ants: Harvard University Press.

Huerta C, Halffter G, Fresneau D. 1992. Inhibition of stridulation in Nicrophorus (Coleoptera: Silphidae): consequences for reproduction. Elytron:151–157.

Kavelaars MM, Lens L, Müller W. 2019. Sharing the burden: on the division of parental care and vocalizations during incubation. Behavioral Ecology. 30:1062–1068.

Korb J, Buschmann M, Schafberg S, Liebig J, Bagnères A-G. 2012. Brood care and social evolution in termites. Proc Biol Sci. 279:2662–2671.

Kramer J, Meunier J. 2018. The other facets of family life and their role in the evolution of animal sociality. Biol Rev Camb Philos Soc.

Lambert GA, Smiseth PT. 2024. Flexible females: nutritional state influences biparental cooperation in a burying beetle. Behavioral Ecology. 35:arae009.

Lucass C, Eens M, Müller W. 2016. When ambient noise impairs parent-offspring communication. Environ Pollut. 212:592–597.

Magrath RD, Haff TM, Horn AG, Leonard ML. 2010. Calling in the Face of Danger. In: Brockmann JH, Roper TJ, Naguib M, Wynne-Edwards KE, Mitani JC, Simmons LW, editors. Volume 41: Elsevier. p. 187–253.

Mariette MM. 2019. Acoustic Cooperation: Acoustic Communication Regulates Conflict and Cooperation Within the Family. Front. Ecol. Evol. 7.

Mariette MM, Buchanan KL. 2016. Prenatal acoustic communication programs offspring for high posthatching temperatures in a songbird. Science. 353:812–814.

Mariette MM, Griffith SC. 2012. Nest visit synchrony is high and correlates with reproductive success in the wild Zebra finch Taeniopygia guttata. Journal of Avian Biology. 43:131–140.

Mariette MM, Griffith SC. 2015. The adaptive significance of provisioning and foraging coordination between breeding partners. Am Nat. 185:270–280.

Moss JB, Tumulty JP, Fischer EK. 2023. Evolution of acoustic signals associated with cooperative parental behavior in a poison frog. Proc Natl Acad Sci U S A. 120:e2218956120.

Müller JK, Eggert A-K. 1990. Time-dependent shifts between infanticidal and parental behavior in female burying beetles a mechanism of indirect mother-offspring recognition. Behavioral Ecology. 27:11–16.

Niemitz C. 1972. Bioakustische, verhaltensphysiologische und morphologische Untersuchungen an Necrophorus vespillo (Fab.). Forma et functio. An international journal of functional biology. 5:209–230.

Niemitz C, Krampe A. 1971. Gehörsinn bei polyphagen Käfern nachgewiesen. Naturwissenschaften. 58:368–369.

Niemitz C, Krampe A. 1972. Untersuchungen zum Orientierungsverhalten der Larven von Necrophorus vespillo F. (Silphidae Coleoptera). Zeitschrift für Tierpsychologie. 30:456–463.

Phillips ME, Chio G, Hall CL, Hofstede HM ter, Howard DR. Seismic noise influences brood size dynamics in a subterranean insect with biparental care.

Potticary AL, Belk MC, Creighton JC, Ito M, Kilner R, Komdeur J, Royle NJ, Rubenstein DR, Schrader M, Shen S-F, Sikes DS, Smiseth PT, Smith R, Steiger S, Trumbo ST, Moore AJ. 2024. Revisiting the ecology and evolution of burying beetle behavior (Staphylinidae: Silphinae). Ecol Evol. 14:e70175.

Prang MA, Zywucki L, Körner M, Steiger S. 2022. Differences in sibling cooperation in presence and absence of parental care in a genus with interspecific variation in offspring dependence. Evolution. 76:320–331.

Pukowski E. 1933. Ökologische Untersuchungen an Necrophorus f. Zeitschrift für Morphologie und Ökologie der Tiere. 27:518–586.

Rauter CM, Moore AJ. 1999. Do honest signalling models of offspring solicitation apply to insects? Proceedings of the Royal Society B: Biological Sciences. 266:1691–1696.

Royle NJ, Smiseth PT, Kolliker M, editors. 2012a. The evolution of parental care. Oxford: Oxford University Press.

Royle NJ, Smiseth PT, Kolliker M. 2012b. The evolution of parental care. Oxford: Oxford University Press.

Rumpf M, Tzschentke B. 2010. Perinatal Acoustic Communication in Birds: Why Do Birds Vocalize in the Egg? The Open Ornithology Journal. 3:141–149.

Sahm J, Brobeil B, Grubmüller E, Conrad T, Schott M, Stökl J, Steiger S. 2024. The scent of offspring: chemical profiles of larvae change during development and affect parental behavior in a burying beetle. Behav Ecol. 35:arae061.

Schrader M, Galanek J. 2022. Stridulation is unimportant for effective parental care in two species of burying beetle. Ecological Entomology. 47:18–24.

Schumacher R. 1973. Beitrag zur Kenntnis der Stridulationsapparate einheimischer Necrophorus-Arten (Necrophorus humator Ol., Necrophorus investigator Zetterst., Necrophorus vespilloides Herbst) (Insecta, Coleoptera). Z. Morph. Tiere. 75:65–75.

Scott MP. 1998. The ecology and behavior of burying beetles. Annu Rev Entomol. 43:595– 618.

Shreeves G, Field J. 2007. Parental care and sexual size dimorphism in wasps and bees. Behavioral Ecology Sociobiology. 62:843–852.

Smiseth PT, Darwell CT, Moore AJ. 2003. Partial begging: an empirical model for the early evolution of offspring signalling. Proc Biol Sci. 270:1773–1777.

Smiseth PT, Moore AJ. 2004. Behavioral dynamics between caring males and females in a beetle with facultative biparental care. Behavioral Ecology. 15:621–628.

Szathmáry E, Maynard Smith J. 1995. The major evolutionary transitions. Nature:227–232.

Trivers RL. 2017. Parental Investment and Sexual Selection. In: Trivers RL, editor. Sexual Selection and the Descent of Man: Routledge. p. 136–179.

